# Phase delays between mouse globus pallidus neurons entrained by common oscillatory drive arise from their intrinsic properties, not their coupling

**DOI:** 10.1101/2024.02.19.580929

**Authors:** Erick Olivares, Charles J. Wilson, Joshua A. Goldberg

## Abstract

A hallmark of Parkinson’s disease is the appearance of correlated oscillatory discharge throughout the cortico-basal ganglia (BG) circuits. In the primate globus pallidus (GP), where the discharge of GP neurons is normally uncorrelated, pairs of GP neurons exhibit oscillatory spike correlations with a broad distribution of pairwise phase delays in experimental parkinsonism. The transition to oscillatory correlations is thought to indicate the collapse of the normally segregated information channels traversing the BG. The large phase delays are thought to reflect pathological changes in synaptic connectivity in the BG. Here we study the structure and phase delays of spike correlations measured from neurons in the mouse external GP (GPe) subjected to identical 1-100 Hz sinusoidal drive but recorded in separate experiments. First, we find that spectral modes of a GPe neuron’s empirical instantaneous phase response curve (iPRC), elucidate at what phases of the oscillatory drive the GPe neuron locks when it is entrained, and the distribution of phases at which it spikes when it is not. Then, we show that in this case the pairwise spike cross-correlation equals the cross-correlation function of these spike phase distributions. Finally, we show that the distribution of GPe phase delays arises from the diversity of iPRCs, and is broadened when the neurons become entrained. Modeling GPe networks with realistic intranuclear connectivity demonstrates that the connectivity decorrelates GPe neurons without affecting phase delays. Thus, common oscillatory input gives rise to GPe correlations whose structure and pairwise phase delays reflect their intrinsic properties captured by their iPRCs.

**Significance Statement:** The external globus pallidus (GPe) is a hub in the basal ganglia, whose neurons impose a barrage of inhibitory synaptic currents on neurons of the subthalamic nucleus, substantia nigra and internal globus pallidus. GPe neurons normally fire independently, but in experimental parkinsonism, they become correlated in the frequency range associated with the pathological rhythms seen in human Parkinson’s disease, raising the possibility that they may be generators of the pathological oscillation. We drove individual pallidal neurons with an oscillatory input over a wide range of frequencies. Cross-correlations of these neurons reproduced many of the features seen in parkinsonism, suggesting that their correlated oscillations might derive from a shared input rather than internal interconnections.

## Introduction

The external globus pallidus (GPe) is thought to maintain parallel independent channels of information flow via the cortico-basal ganglia (BG) circuit (Alexander et al., 1986; Foster at al., 2021), which are necessary for effective action selection (Mink, 1996; Boraud et al., 2000), despite the strongly convergent innervation of GPe neurons (Yelnik et al., 1984; Shink and Smith, 1995; Oorschot, 1996; Zheng and Wilson, 2002; Bar-Gad et al., 2003b; Kita et al., 2004; Goldberg and Bergman, 2011). Key experimental evidence for the independent processing within the GPe is the observation that the firing of GPe neurons is uncorrelated under normal conditions (Nini et al., 1995; Raz et al., 2000; Goldberg et al., 2002; Heimer et al., 2002; Bar-Gad et al., 2003a; Kita et al., 2004; Mallet et al., 2008; Goldberg and Bergman, 2011). GPe firing is uncorrelated because any two GPe neurons share very little common striatal input. Moreover, the GPe intrinsic connectivity can actively decorrelate GPe firing (Wilson, 2013).

A hallmark of parkinsonian pathophysiology both in animal models (Raz et al., 1996; Goldberg et al., 2002; Goldberg et al., 2004; Mallet et al., 2008) and in humans (Brown, 2006; Hammond et al., 2007; Cagnan et al., 2019; Crompe et al., 2020) is the appearance of beta-range oscillations coherently throughout the cortico-BG circuits. These oscillations are widely viewed as arising from the destabilization of negative feedback loops that are embedded within the cortico-BG circuit (Plotkin and Goldberg, 2019), most prominently the GPe-subthalamic circuit (Plenz and Kital 1999; Terman et al., 2002; Tachibana et al., 2011) following dopamine denervation of the BG. The beta oscillations are visible as sinusoidal fluctuations in the local field potentials (LFPs) recorded in the cortico-BG circuit (Brown and Williams, 2005), including in the GPe (Goldberg et al., 2004). We have shown that LFPs are highly synchronous throughout the GPe of parkinsonian primates (Goldberg et al., 2004). Because the LFP represents the synaptic currents flowing into nearby neurons (Creutzfeld et al., 1966; Elul 1972; Mitzdorf, 1985; Destexhe et al., 1999; Lass, 1986; Goldberg et al., 2004; Haidar et al., 2016), the presence of synchronous oscillations in the GPe LFP of parkinsonian animals, means that pairs of GPe neurons, even several millimeters apart, receive strong common oscillatory synaptic input. Indeed, in experimental parkinsonism, the pairwise cross-correlations between the firing of GPe neurons have a beta-range oscillatory structure (Nini et al., 1995; Raz et al., 2000; Goldberg et al., 2002; Heimer et al., 2002; Heimer et al., 2006; Mallet et al., 2008; Plotkin and Goldberg, 2019).

Neurons receiving common oscillatory input are expected to fire in synchrony with each other (Perkel et al., 1967). However, the oscillatory cross-correlations of GPe neurons recorded in parkinsonian primates reveal large phase delays among these neurons. This curious observation led us to ask what determines the structure of the cross-correlation between GPe neurons driven by common input. In particular, how can GPe neurons receiving common sinusoidal input fire with non-zero phase delays relative to each other; and do these phase delays arise from intrinsic properties of GPe neurons or from their mutual connectivity? GPe neurons are autonomous pacemakers whose firing becomes entrained by oscillatory inputs over a wide frequency range of 10-70 Hz (Chan et al., 2004; Deister et al., 2013; Wilson and Jones, 2023). We therefore addressed the following questions. What determines the distribution of stimulus phases at which each GPe neuron fires when driven by sinusoidal input, and how does this change with the frequency of the input? Can the phase delays among pairs of neurons be predicted from the differences in these spike phase distributions? Can common drive generate a broad range of phase differences even when the two GPe neurons are entirely independent of each other? How might the known intrinsic connectivity among GPe neurons influence the correlations among them both in term of their strength and phase differences?

## Materials and Methods

### Institutional approval

All experiments were conducted in accordance with the National Institutes of Health guidelines and were approved by the Institutional Animal Care and Use Committee of The University of Texas at San Antonio.

### Animals

The animal subjects used in this research have been detailed previously in Wilson & Jones 2023. The study involved 14 mice aged between 74 to 193 days. Out of these, 10 were PV-Cre, 2 were Npas1-Cre-tdTomato mice, and 2 were wildtype C57BL/6 mice without any genetic modifications. In total, 16 GPe neurons were examined, all classified as prototypic cells using the electrophysiological criterion of Jones et al. (2023). Eleven of these were confirmed as parvalbumin-positive (PV+) either through fluorescence or via a distinct light-induced Archaerhodopsin-3 (Arch) hyperpolarization. The remaining neurons were identified in slice preparations that lacked any PV reporter.

### Acute brain slices

Mice were deeply anesthetized with isoflurane and killed by decapitation. Parasagittal brain slices of 300 µm that included the GPe were obtained using a vibrating microtome (Leica) in an ice-cold cutting solution. This solution consisted of the following (in mM): 110 choline chloride, 2.5 KCl, 1.25 NaH_2_PO_4_, 0.5 CaCl_2_, 7 MgSO_4_, 25 glucose, 11.6 Na ascorbate, 3.1 Na pyruvate, and 26 NaHCO_3_, and was saturated with 95% O_2_ and 5% CO_2_. The slices were then transferred to ACSF, which contained (in mM): 126 NaCl, 2.5 KCl, 1.25 NaH_2_PO_4_, 2 CaCl_2_, 1 MgSO_4_, and 10 glucose, also bubbled with 95% O_2_ and 5% CO_2_. The ACSF used for storing the slices, but not for recording, additionally included (in mM): 0.005 glutathione, 1 Na ascorbate, and 1 Na pyruvate. The slices were subsequently heated to 34°C for 30 minutes and then permitted to cool to room temperature before being used.

### Perforated patch recordings

The slices were continuously superfused with ACSF oxygenated and heated to between 33°C and 35°C. Neurons were imaged using an Olympus BX51WI microscope equipped with a 40X water-immersion lens and Dodt gradient contrast optics. Recordings of neurons were made using the perforated patch-clamp method. Recording pipettes made from filamented borosilicate glass (G150F-4; Warner Instruments) were pulled to have resistances of 3-7 MΩ using a Flaming-Brown pipette puller (model P-97; Sutter Instruments). The pipette tip was initially filled with a solution containing (in mM): 140 KMeSO_4_, 10 HEPES, 7.5 NaCl, and then back-filled with the same solution plus 1 *μ*M Gramicidin-D (MP Biomedicals). After achieving a gigaohm seal, a period of 10-30 minutes was allowed for the gramicidin to perforate the cell membrane, ensuring adequate electrical access (40-70 MΩ). Recordings were obtained using a MultiClamp 700B amplifier (Molecular Devices), connected to an ITC-18 A/D converter (HEKA Instruments), and managed on a Macintosh computer with custom software written for Igor Pro (WaveMetrics).

### Oscillatory driving and vector strength

Neurons were driven by sinusoidal currents with frequencies ranging from 1 to 100 Hz in 1 Hz steps. Each frequency was presented for 10 s. Spike times *t*_*j*_ were converted to phases using *ϕ*_*j*_ = *ft*_*j*_ mod 1 where *f* is the driving frequency. To assess spike entrainment to the stimulus at each frequency, we calculated the vector strength (VS) as:

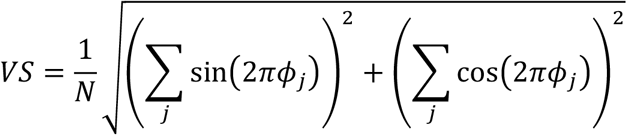

where *N* is the number of spikes. Additionally, the vector angle (VA) was computed as:

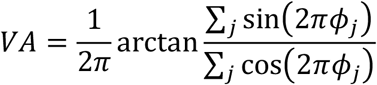

The VA provides an estimate of the average phase on the input oscillation at which the neuron fired, while the VS indicates the reliability of firing at that angle. A vector strength of zero signifies firing at random with respect to the oscillation, while a value of one denotes firing consistently at the same exact phase on the oscillation on every cycle.

### iPRC measurements

The infinitesimal PRC was estimated as described by Wilson et al. (2014). The intrinsic oscillation of the neuron was perturbed by a barrage of contiguous brief pulses of injected current (0.25 ms) with amplitudes drawn from a Gaussian distribution (mean = 0, standard deviation (SD) = 40 pA). A total of 40 episodes, each lasting 4 s, were applied to each neuron. Each inter-spike interval (ISI) was divided into 50 equal-length bins, and the charge applied in each bin was calculated. The resulting matrix of applied charges was subjected to multiple regression analysis to estimate the iPRC values at each bin.

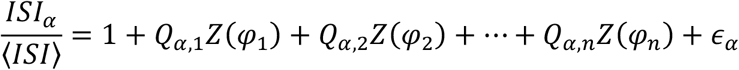

The resulting iPRCs were parameterized as in Olivares et al. (2022) using an *ad hoc* function with 7 free parameters that has proven capable of capturing the shape and size of the GPe iPRC. The parameterized iPRC was used to simulate the phase equation (see below) and obtain the iPRC Fourier components.

### Representation of the iPRC as a Fourier series

The autonomous pacemaking of a GPe neuron is described as an oscillator whose intrinsic phase is *φ ∈* [0,1) and whose intrinsic frequency is *f*. Consequently, the neuron’s iPRC is also a periodic function of phase. According to Fourier theory, any periodic function can be represented as an infinite series of cosine functions with integer frequencies plus a constant term. Additionally, the cosine function of frequency *n*, which is also called the *n*^*th*^ Fourier *mode*, is shifted by a specific angle, which we denote *Δ*_*n*_. Thus, we have (Goldberg et al., 2013; Morales et al., 2020).

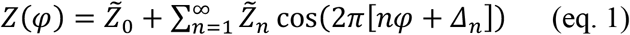

The formulas for the prefactors 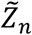 are: 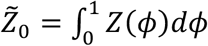 (the temporal average of the iPRC) and 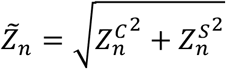 where 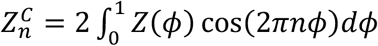 and 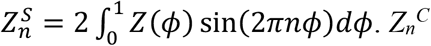 and 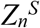 are called the *n*^*th*^ cosine and sine Fourier transforms of *Z*(*φ*), and are essentially the correlation between the iPRC and *n*^*th*^ cosine and sine functions, respectively. The angles are given by the formula 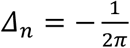 arctan 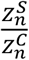 On a practical level, summing a finite number of modes in the above series (e.g., beginning with modes 0,1,2, and so on) up to some *N*^*th*^ mode results in an ever-improving approximation of the iPRC as *N* is increased.

### Relationship between the phase of the iPRC modes and the stimulus phase of locking

In order to gain insight into the phase at which a GPe neurons will phase lock to an oscillatory drive, it is informative to consider how the phase oscillator representing the GPe neuron is affected by sinusoidal driving at the same intrinsic frequency *f* of the oscillator (or integer *n* multiples of *f*). In this case, the dynamics of the phase oscillator are governed by the following phase equation

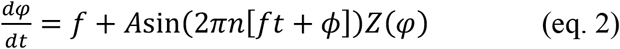

Because the oscillator is being driven at its intrinsic frequency, it stands to reason that the oscillator will continue to oscillate at its intrinsic frequency when driven, so we use the approximation that phase advances linearly, i.e., 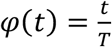, where *T* (*= 1/f*) is the period of the oscillator. In this approximation, we can now integrate the phase equation (eq. 2) between *0* and *T*. In the case of phase locking at the intrinsic frequency the sinusoidal driving should add no net phase change when averaged over the neuron’s period, so that the neuron will spike at the same phase of the input on every period of driving. In other words, we are looking for a stimulus phase *ψ* such that

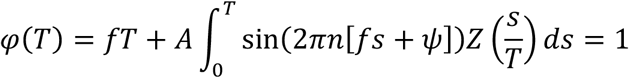

Because *fT = 1* we have

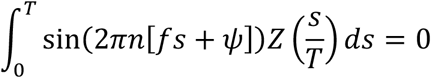

While this equation for *ψ* can be solved directly, it is instructive to expand the left-hand-side of this equation using Fourier representation of the iPRC (eq. 1).

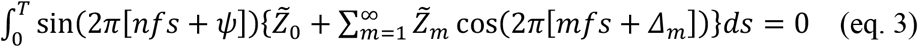

The first term in eq. 3 vanishes because the integral of a sinusoidal function over one period is zero (multiplication by the constant 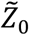 does not change that). The rest of the infinite number of terms in eq. 3 all have the following structure

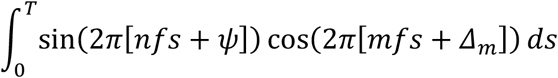

Using trigonometric identities (plus, again, the useful result that the integral of a sinusoidal function over one period vanishes) demonstrates that for all iPRC modes for which *m ≠ n* the integral vanishes. Only when *m = n*, does the integral equal sin(2*π*[*ψ – Δ*_*n*_]) (up to a scale factor). This underscores the finding that when you drive an oscillator at an integer multiple (*n*), or harmonic, of its fundamental frequency you are only affecting (or interacting with) the corresponding *n*^*th*^ mode of its iPRC (Goldberg et al., 2013). All other modes are unaffected, and are said to be *orthogonal* to the input. Additionally, all Fourier modes are orthogonal to each other because

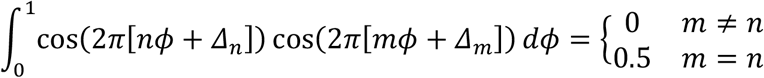

Thus, eq. 3 reduces to sin(2*π*[*ψ − Δ*_*n*_]) *=* 0, or in other words, *ψ = Δ*_*n*_. The meaning of this result is that the sine wave perturbation needs to be shifted exactly by the angle of the *n*^*th*^ mode of the iPRC when driven by *n* times the fundamental frequency in order to create zero phase perturbation and therefore stable locking (the other solution *ψ =* 0.5 + *Δ*_*n*_ is unstable). In other words, even when the phase oscillator is driven at precisely one of its harmonic frequencies, there are specific phases at which the input would still be orthogonal to the corresponding mode and not drive any phase change.

### Parameterization of a triangular iPRC

It is instructive to consider the triangular iPRC (of unity height) that is parameterized by the location *θ* of its peak [also because it is a good first approximation for some empirical iPRCs (Tiroshi and Goldberg, 2019)]. The sine and cosine Fourier modes of the triangular iPRC are given by 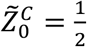 and for all *n > 0* by

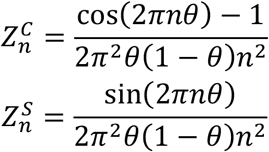

In this model, a straightforward calculation shows that the angle of the iPRC’s fundamental frequency (and hence the phase at which an oscillator with the same frequency will stably lock) is 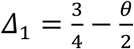

### Numerical integration of the phase equation and additive noise

Neurons’ return maps (RMap) were obtained by numerically integrating the phase model equation (eq. 1) using Euler’s method with a constant time step of 0.05 ms. For each stimulus frequency *f*, we performed 400 independent simulations starting at equidistant stimulus phases (relative to a preceding spike) in the range [0, 1). For each starting phase *ϕ*_*prev*_, a deterministic simulation was initialized with the neuron phase *φ = 0*. The stimulus was run until the neuron spiked (*φ = 1*), and the stimulus phase at the moment of the spike was recorded as *ϕ*_*next*_. The return map is fully determined by the neuron’s iPRC, intrinsic rate, and the stimulus frequency and amplitude.

We used the calculated return map to obtain a sample of neurons’ spike phases. The sample was obtained by iterations in the return map, discarding the first 100 iterations. In every iteration, additive noise was introduced to account for the neurons’ intrinsic spike variability. The iteration rule was defined as:

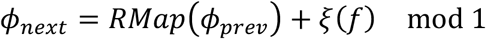

Where *ξ*(*f*) is additive Gaussian noise that depends on the stimulus frequency *f* with zero mean and SD of 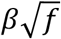. *β* was tuned to match the spread of spike phases when the neuron is driven by a stimulus frequency closer to the neuron’s intrinsic spike rate (i.e., when locking occurs). We adjusted the standard deviation in proportion to the square root of the stimulation frequency. This adjustment was necessary because as the frequency increases, the stimuli period becomes shorter, making the return map predictions less precise.

### GPe neural network model

The GPe neural network was simulated as described in Olivares et al. (2022 One thousand model neurons were connected using a small-world architecture; each neuron was connected in a ring-like organization to its ten closest neighbors, and then 1% of the synapses were randomly rewired. The degree of connectivity is consistent with experimental data (Sadek et al., 2007; Bugaysen et al., 2013; Higgs et al., 2021, Jones et al., 2023). Neurons in the network were heterogeneous with respect to their iPRCs and intrinsic firing rates; both neuron properties were drawn from distributions that qualitatively represent experimental data from GPe neurons. The unitary inhibitory synaptic conductance for the local synaptic connections was modeled with an instantaneous rise and an exponential decay with a time constant of 2.4 ms. The peak amplitude of each local synaptic conductance varied independently from spike to spike to represent quantal synaptic fluctuations and was drawn from a normal distribution with a mean of 2.5 nS and a SD of 1.75 nS. The reversal potential was set at -73 mV. These values were chosen to match those of spontaneous unitary synaptic conductances observed in voltage clamp recordings from GPe neurons (Higgs et al., 2021).

When unconnected, the network was simulated with all synapses removed, and the neurons were driven solely by the sinusoidal input (20 pA amplitude) and an additive noise tuned to each cell using the theoretical approximation in (Ermentrout et al., 2011) to generate an inter-spike interval coefficient of variation of 0.08. When connected, we compensated for the decrease in rate due to the local inhibitory inputs using an excitatory direct current tuned to each neuron individually. To discriminate the effect of connectivity from the effect of the barrage of inhibitory synaptic inputs produced by the local network in the absence of external drive, we simulate a disconnected network in which every neuron received a barrage of synaptic inputs with a mean rate equal to the one received in the connected configuration, but without the temporal structure produced by synaptic interactions and sinusoidal drive. To evaluate the effect of connectivity pattern *per se*, we repeated the simulations using either random or rich-club architectures (long-tailed distribution in the number of postsynaptic connections per neuron), while maintaining the number of connections. To evaluate the effect of the degree of connectivity, we evaluated the network using double and half of the sparseness estimated from the literature (20 and 5 incoming synapses in average, respectively).

Network simulations were conducted using a C++ package originally designed for continuous-time recurrent neural networks (Beer, 1995), accessible at https://rdbeer.pages.iu.edu. We made modifications to the package, introducing the phase model and incorporating our synaptic events. The model was numerically integrated using Euler’s method, employing a fixed time step of 0.1 ms. Common external stimuli were applied, covering frequencies from 1 to 100 Hz, and each frequency was simulated for a duration of 100 seconds.

### Deriving the cross-intensity function (CIF) from the spike phase distribution of neurons driven by a common sinusoidal input

Assume we have two spike trains that are driven by common sinusoidal input of frequency *f*

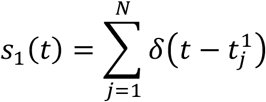

and

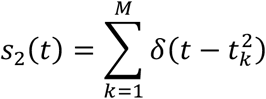

where 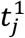 and 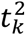 are the spike times of the two neurons, respectively.

The cross-intensity function (CIF), which is the spike cross-correlation function (Cox and Lewis, 1966; Perkel et al., 1967), is given by

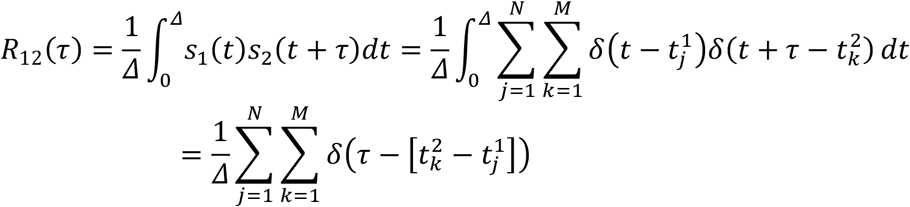

If we restrict ourselves to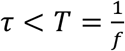, then the only pairs of spikes 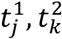 that contribute to the sum are those that appear in the same period of the sinusoidal input. But in this case, we can define probability distributions *P*_1_(*ϕ; f*) and *P*_2_(*ϕ; f*) of the phase at which the spikes 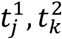 occur, which we will refer to as the *spike phase distributions*, and use the ergodic theorem to replace the time average (*1/Δ*) by averages on these distributions of the Dirac delta function. This yields

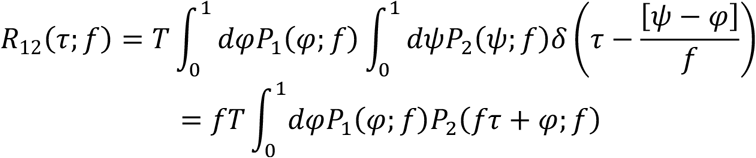

where the pre-factor *T* is because we averaged over time Δ for the spikes of the first neuron and averaged a neighborhood of *T* for the spikes of the second (so we need to multiply back by *T*). Then according to the algebra of the Dirac delta function we need to multiply times *f*. But because *fT = 1* we get

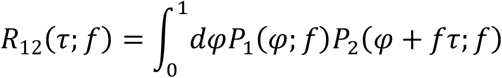

This means that the CIF of spikes between *0* and *T* is equal to the cross-correlation function of the probability distribution functions.

### Statistics

We used Pearson correlation (*r*) to measure the linear association between variables, and Wald’s test to determine the statistical significance of the association. Calculations were done using SciPy.Stats Python library. The null hypothesis of no linear association was rejected if the P value was smaller than 0.05.

## Results

### GPe neurons driven periodically exhibit phase locking

GPe neurons (*n = 16*) were recorded in acute brain slices (from *N = 14* mice) in the perforated patch configuration to preserve their autonomous firing over the long recording duration. Each neuron was subjected to a sequence of 100 ten-second-long sinusoidal current waveforms at frequencies 1, 2, 3, …, 100 Hz, each with an amplitude of 20 pA. This amplitude, although small in comparison to *in vivo* synaptic currents, proved to be sufficient for inducing rate modulation and spike entrainment in our experimental setup. In 3 of the 16 neurons the cell did not survive all 100 trials, but all neurons were recorded up to 50 Hz. As we reported previously (Wilson and Jones 2023), the sinusoidal drive influences the timing of the spikes (Fig. 1A) causing the spike times to distribute non-uniformly across the phases of the sinusoidal waveform in a frequency-dependent manner. We call this distribution the *spike phase distribution* and denote it *P*_*j*_(*ϕ; f*) of the *j*^*th*^ neuron for driving frequency *f*. In the example depicted in Fig. 1B, for low and high driving frequencies (top and bottom panels), the GPe spikes distributed broadly across phases – the spike phase distribution itself was sinusoidal – and peaked at phase 0.25, which corresponds to the peak of the sinusoidal waveform. However, when the driving frequency was close to the intrinsic firing rate of the neuron, the spike phase distribution became narrower and peaked at a later phase that varied among neurons (approximately 0.4 in Fig. 1B, middle). The degree of phase locking is captured by the vector strength (VS, Fig. 1D, see Materials and Methods), which increases as the distribution in Figure 1B is more sharply peaked (and unimodal). The VS increased as the driving frequency approached the intrinsic firing rate of the neuron, indicating that the neuron was more strongly entrained when driven by a frequency similar to its own intrinsic rate. As the driving frequency approached the intrinsic rate of the GPe neurons, the cyclic averaged phase angle of firing (VA, Fig. 1E) shifted away from stimulus phase 0.25 (the peak of the sinusoidal driving current) towards larger phase shifts. What determines the stimulus phase at which the GPe neurons became maximally entrained, and why does this phase vary among neurons?

**Figure 1.**
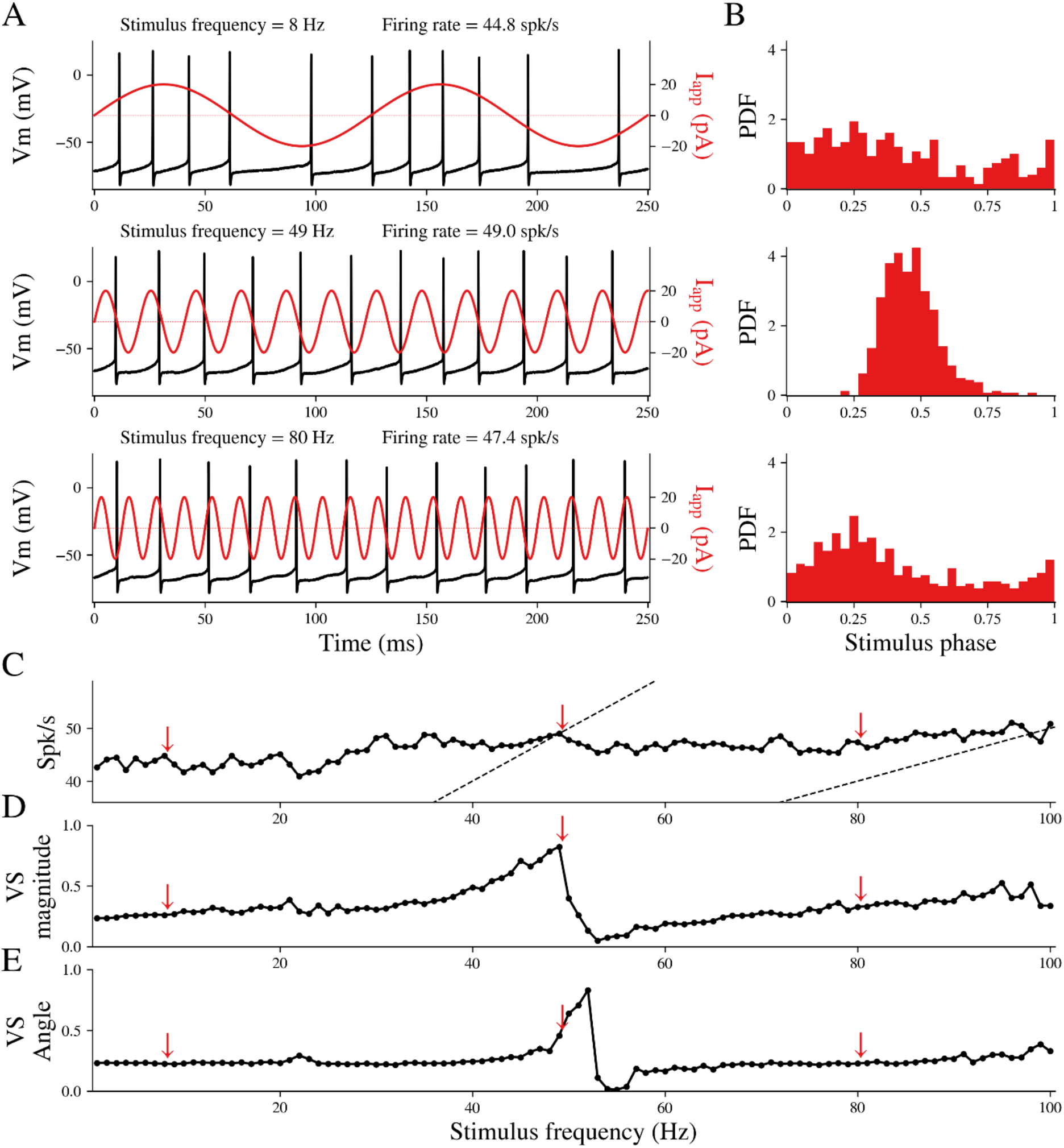
GPe neurons driven periodically exhibit phase locking. A) Voltage trace of one example cell driven by a periodic current input at three different frequencies. Spike times were mapped to stimulus phase, and the distributions of spike phases are shown in B. C) Mean firing rate of the neuron during the 100 periods of stimulation. Red arrows indicate the stimulation frequencies shown in A and B. Dashed lines indicate a 1:1 ratio of firing rate to driving frequency (left) and a 1:2 ratio (right). D) Vector strength and E) vector angle as a function of stimulus frequency for the example cell.

### Fourier modes of the iPRC dictate the phase at which phase locking occurs

We hypothesized that the diversity of stimulus phases at which different GPe neurons entrain is related to the heterogeneity of the their iPRCs. To see why, we first digress with a simplified iPRC. Consider a triangular iPRC (of unity height) that is driven by a sinusoidal input whose frequency matches the neuron’s intrinsic firing rate. The iPRC is parameterized by a single parameter *θ*, which is the location (between 0 and 1) at which the triangle peaks (Fig. 2A). As described in the Materials and Methods section, the iPRC can be decomposed into (a constant term plus) an infinite series of cosine functions each with: 1) a frequency that is an integer (*n = 1,2,3*…) multiple (or harmonic) of the fundamental frequency; 2) an amplitude; and c) and angle, between 0 and 1 (eq. 1 in Materials and Methods). Figure 2B depicts the first four cosine functions, or *modes*, of the triangular iPRC. Their amplitude is indicated with a black vertical line and their angle is the distance of that line from 1 (indicated by a red line) *divided* by the period of that mode (indicated by a magenta line). The amplitudes of the modes decrease like *1/n*^*2*^ (see Materials and Methods) and their angles increase linearly (Fig. 2C). Figure 2A includes a sequence of curves, each representing a partial sum of the first *N* terms (with *N = 0, 1, 2, 3, 4*). As the number of modes used in the partial sum is increased the approximation of the iPRC improves.

**Figure 2.**
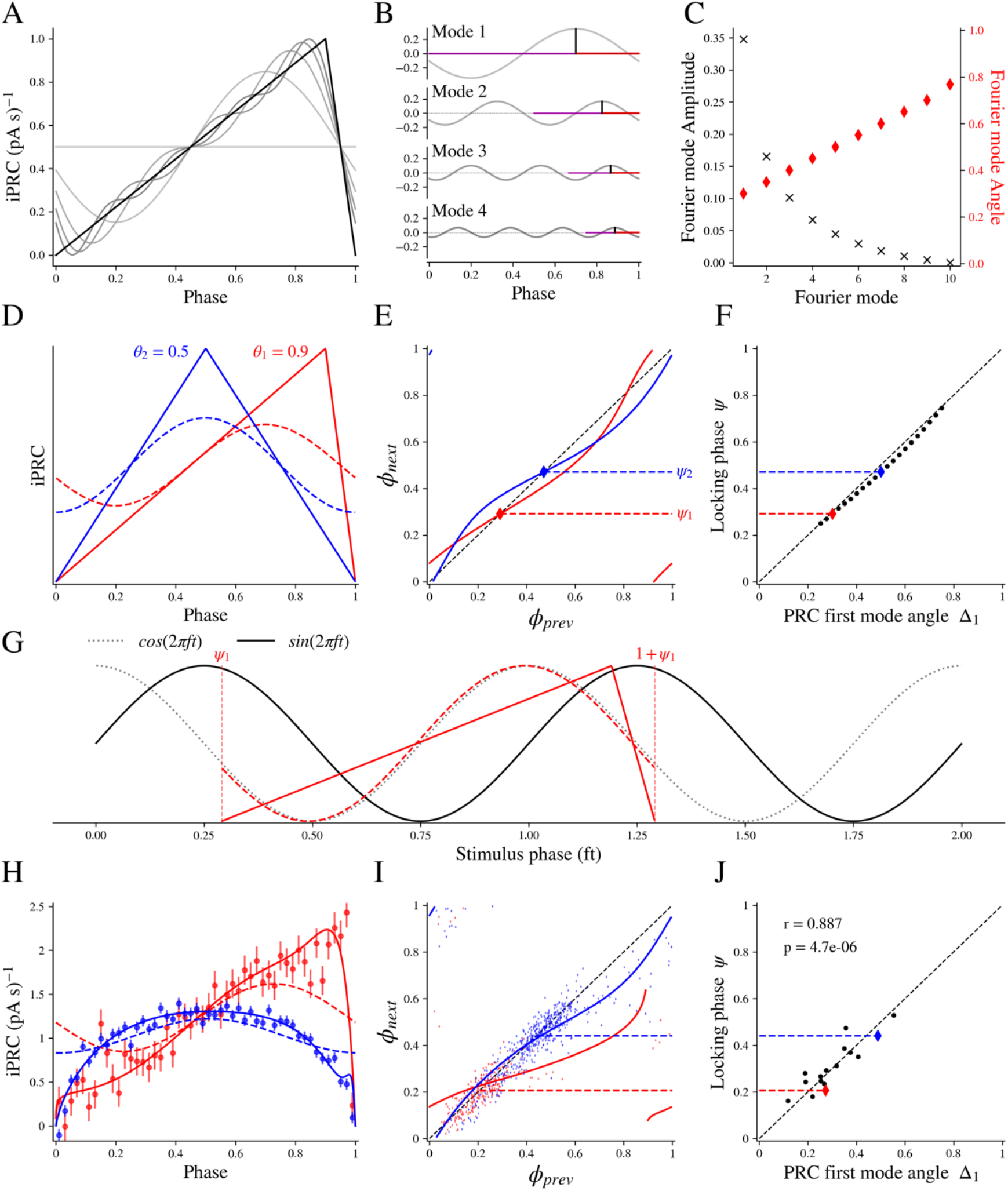
Fourier modes of the iPRC dictate the phase of spikes on the sine wave input at locking. A) Triangular iPRCs with *θ* value of (i.e., location of peak at) 0.9 (solid black) and sequences of curves representing a partial sum of the first *N* terms or modes of the Fourier decomposition (with *N = 0, 1, 2, 3, 4*, gray lines). B) First four modes of the iPRC shown in A, black vertical line show the mode amplitude and their angle is the represented by the fraction of the horizontal red line in proportion to the period of that mode, indicated by a magenta line. C) Amplitudes (black exes) and angles (red diamonds) of the first 10 modes of the iPRC shown in A. E) Triangular iPRCs with *θ* values of 0.5 (blue) and 0.9 (red), dashed lines shows the first Fourier mode component of each iPRCs. F) Predicted return map indicating the phases of sequential spikes when each of the iPRCs shown in A is driven by a sinusoidal stimulus at the neuron’s natural frequency; diamonds and dashed lines show the stable locking phase (ψ) for each iPRC. F) Phase at locking (ψ) closely matches the Fourier first mode of the iPRC (Δ1) as predicted by the theoretical approximation. G) Relationship between the sinusoidal input (black solid line), the iPRC (solid colored line), its first Fourier mode (dashed colored line), and the orthogonal function to the stimulus (cosine, dashed black line). The iPRC and its first Fourier mode are aligned with the stimulus phase at two sequential spike times (ψ & 1 + ψ, vertical lines). It is apparent that the Fourier mode of the iPRC nearly merges with the cosine function orthogonal to the sine wave stimulus. All curves are normalized for easy comparison. H) Two examples of iPRCs estimated from GPe neurons; filled symbols and error bars show the iPRC estimated by the multilinear regression method, solid lines show the parameterized fit to each iPRC and dashed lines shows the first Fourier mode component of the iPRCs. I) Solid lines show return maps predicted by the phase model. Colored dots show *ϕ*_*next*_ vs. *ϕ*_*prev*_ in the experimental data. Locking phases indicated by horizontal dashed lines are calculated as the vector angle. J) Correspondence between the phase of the first mode of the iPRC and the phase of locking, calculated as the vector angle for the spikes at the locking frequency for all 16 cells in the sample (r = 0.887, P < 10^-5^).

In order to calculate the phase of the sinusoidal input at which the neuron will lock to its sinusoidal input, we will consider two different triangular iPRCs, a skewed one (*θ =* 0.9) and a symmetrical one (*θ =* 0.5) (Fig. 2D). We need to first calculate the return map that represents the phase of the next spike on the input, *ϕ*_*next*_, as a function of the previous spike’s phase, *ϕ*_*prev*_ (Fig. 2E). This is achieved by integrating the phase equation of the neuron driven by sinusoidal input (eq. 2 in Materials and Methods) for all values of *ϕ*_*prev*_ between 0 and 1, and plotting the resultant *ϕ*_*next*_ (e.g. Wilson, 2017). The phase locking occurs at the phase at which the return map crosses the identity line (diagonal in Fig. 2E) with a slope below 1 (that guarantees that this fixed point on the return map is stable). Note that the stimulus phase of locking depends on the shape of the iPRC. For the rightward skewed iPRC (red) the phase locking occurs slightly above 0.25 and for the symmetrical iPRC (blue) it occurs at approximately 0.5 (Fig. 2E).

To gain an intuitive understanding of what determines the phase locking, we sought an approximate solution of the phase equation (eq. 2 in Materials and Methods) under the assumption that the phase *φ* that appears as the argument of the iPRC [*Z*(*φ*)] can be replaced with the solution of the unperturbed phase equation (namely that 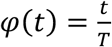). Although this is never strictly true because it assumes the stimulus does not alter cell phase, it is a useful approximation when the stimulus amplitude is small and the driving is at precisely the intrinsic frequency of the neuron (so the period is not expected to change). In this approximation, the phase locking occurs at the phase *ψ* that equals the phase *Δ*_1_of the fundamental mode of the iPRC when it is written as cos(2*π*[*φ* + *Δ*_1_]). For the triangular iPRC, we found (see Materials and Methods) that 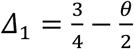. Indeed, if we plot the phase *ψ* at which phase-locking occurs based on the numerical solution of the full phase equation, against *Δ*_1_, we find that the points lie very close to the identity line (Fig. 2F). Fig. 2G clarifies why this is. When driving the neuron with a sinusoid described by sin 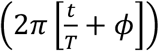 integration of its product with the term 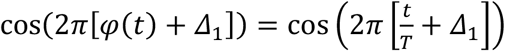 (in the approximated solution of eq. 2) equals zero if *ϕ = Δ*_1_ (because sine and cosine functions of the same frequency are orthogonal if their phase shifts equal each other). Geometrically, this means that if we plot the iPRC so that it spans from one phase-locked spike to the next (vertical dashed lines in Fig. 2G) along the sine wave input (solid black sinusoid in Fig. 2G), then the fundamental Fourier mode of the iPRC (dashed sinusoids in Fig. 2G) will very nearly merge with the cosine wave (dotted black sinusoid in Fig. 2G) that corresponds (i.e., that is shifted to the left by 0.25 relative) to that sine wave. In the approximate solution, where *ψ = Δ*_1_, the fundamental mode of the iPRC merges precisely with the cosine wave. The discrepancy between the approximate and full solutions of the phase equation depends on the shape of the PRC (compare red and blue dots in Fig. 2F, see below) and on the amplitude of the oscillatory driving (which determines to what extent the dotted solutions bulge away from the unity line in Fig. 2F).

Note that in this simplified model the more skewed red empirical iPRC leads to phase locking at a smaller phase shift than the more symmetrical blue empirical iPRC. Thus, the fundamental Fourier mode of the iPRC generates an “equivalence class” among neurons, in the sense that neurons whose iPRCs’ fundamental modes shares the same angle will all lock at the same phase when driven at a frequency equal to their intrinsic firing rate. Conversely, any difference between neurons in the fine structure of their iPRCs that does not affect the fundamental mode will not impact this entrainment and phase locking. Another corollary of this finding is that if the neuron is driven at the *n*^*th*^ harmonic of its intrinsic rate, then the phase locking will occur at the phase of the *n*^*th*^ Fourier mode of the iPRC (see Materials and Methods). In line with a previous study of ours (Goldberg et al., 2013), these findings underscore the importance of the iPRCs’ Fourier modes in understanding how pacemaker neurons respond to periodic inputs (see Discussion).

These results were based on a highly idealized iPRC, and on the 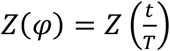 approximation, which is valid only for vanishingly small stimuli (see Materials and Methods). To determine whether the results apply more broadly, we tested the results using the empirical iPRCs of all 16 of our recorded neurons (Fig 2H), and numerically integrated the phase model of each neuron to generate return maps (Fig. 2I). These reproduced the correspondence between the phase *Δ*_1_ of the first mode of the iPRC and the stimulus phase *ψ* of locking, calculated as the vector angle for the neurons’ spikes at the locking frequency (Fig. 2J, Pearson correlation *r = 0*.*887, P < 10*^*-5*^). Note the similarity between the phase of the fundamental modes of the empirical iPRCs (dotted lines in Fig. 2H) and that of the triangular iPRCs (dotted lines in Fig. 2D), which demonstrates the notion of the “equivalence class” of the skewed iPRCs and of the symmetrical ones, and clarifies that it is the fundamental mode that mostly influences the shape of the return maps and their stable fixed points.

### Return maps determine the shape of the spike phase distributions across frequencies

The formalism of using the return maps (which are calculated from the phase equation and using the GPe neurons’ empirical iPRC) to predict the spike phase distributions *P*_*j*_(*ϕ; f*) is not restricted to the case of phase locking. In Fig. 3A, we depict (in red) the scatter plot of all the empirical pairs (*ϕ*_*prev*_, *ϕ*_*next*_) representing consecutive phases of the sinusoidal input at which the GPe neuron spiked. In Fig. 3B, we depict the spike phase histograms (red), which correspond to the density of points on the scatter plot for each frequency. The deterministic return maps predicted from the empirical iPRC for the same three frequencies (Fig. 3A, blue) correspond closely to the red scatter plots. Moreover, if we assume a certain degree of additive noise to the deterministic return maps (see Materials and Methods for how the noise level was determined) than we find that the resulting spike phase distributions (black lines in Fig. 3B) match the empirical histograms (red bars in Fig. 3B). To estimate the goodness-of-fit, we calculated the average across all GPe neurons of the Pearson correlations between the predicted and empirical phase distribution (Fig. B, blue steps vs. red bars, respectively) as a function of the driving frequency (Fig. 3C) and found a high degree of correlation of approximately 0.75 across all frequencies.

**Figure 3.**
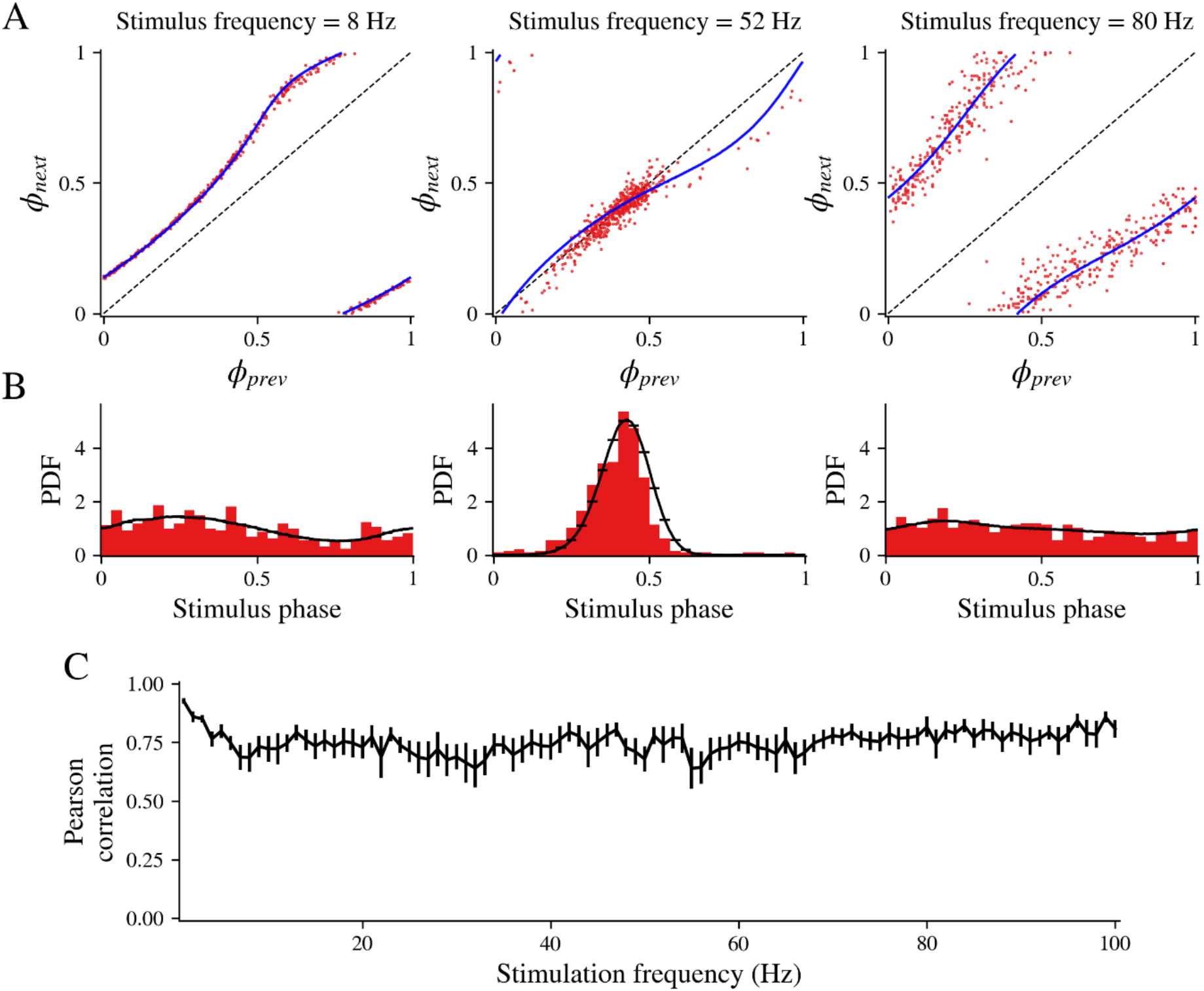
Return maps determine the shape of the spike phase distributions across frequencies. A) Examples illustrating the alignment between a neuron’s return maps and its spike phases at three different frequencies of sinusoidal driving input (8, 52 and 80 Hz). The blue line shows the return map generated by the phase model using the neuron’s iPRC, while red dots depict recorded pairs of consecutive spike phases on the sinusoidal input. B) The spike phase histograms (red) correspond to the density of red points in A. The black line in B shows the predicted density from the return maps for the same three frequencies. C) Pearson correlations between the predicted (black steps) and empirical phase distributions (red bars) were calculated across all GPe neurons and plotted as a function of the driving frequency.

### Pairwise CIFs during sinusoidal driving are predicted from the phase distributions across all frequencies

We have seen that the iPRCs and phase maps can give rise to diverse spike phase distributions, and the modes of these distributions can occur at various phase delays in a frequency-dependent fashion. Can the heterogeneity of iPRC shape among neurons give rise to the diversity of their pairwise CIFs? Intuitively this seems correct. Consider two GPe neurons whose spike phase distributions (at a specific frequency) possess modes (e.g., where they peak) at different phases. The maximal value of their pairwise CIF should be located at the phase difference between these respective modes. In the Materials and Methods, we present the general derivation that shows that the structure of the pairwise CIF of two neurons is equal to the cross-correlation function of their respective spike phase distributions. In Figs. 4A-C, we depict several examples of empirical pairwise CIFs (green curve) with the CIF that is predicted from the two empirical spike phase distributions (purple curve) that are shown to the left (blue and orange distributions). The correspondence is impressive, with even some of the fine structure of the CIF being explained by cross-correlating the phase distributions. To estimate the goodness-of-fit between the predicted and empirical CIFs, we calculated the average across all (*16×15/2* =) 120 GPe neuron pairs of the Pearson correlations between the predicted and empirical CIFs (purple vs. green, respectively) as a function of the driving frequency (Fig. 4D) and found a high degree of correlation of approximately 0.6 across all frequencies. Moreover, because the Pearson correlations are independent of amplitude, we also determined that the amplitudes of the empirical CIFs were highly correlated the amplitude of predicted CIFs (Fig. 4E).

**Figure 4.**
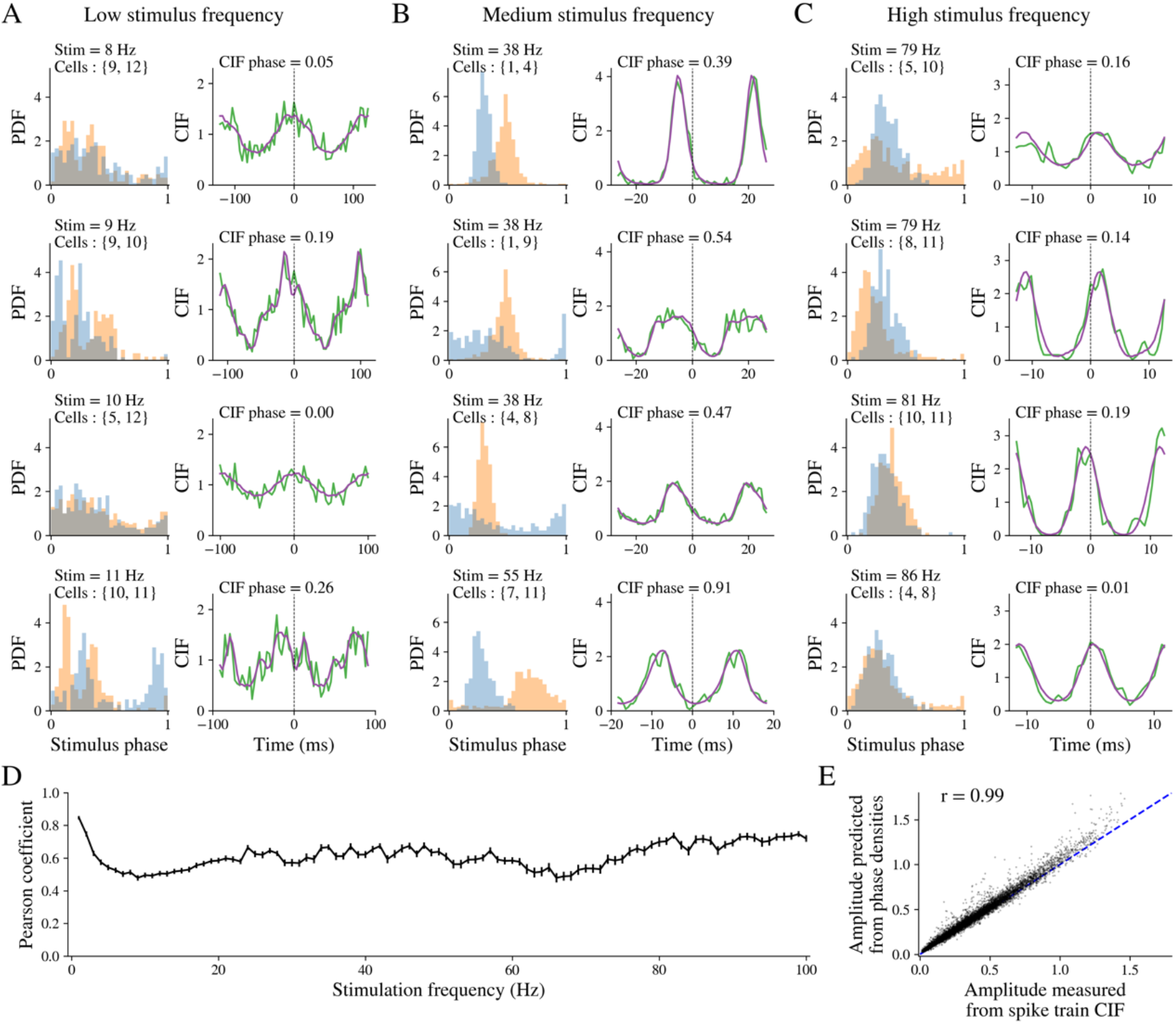
Pairwise CIFs during sinusoidal driving are predicted from the spike phase distributions across all frequencies. A-C) Left: Examples of spike phase distribution functions (PDFs) for two neurons (blue and orange), with cell pairs and stimulus frequencies shown in the plots. Right: Corresponding empirical pairwise cross-intensity functions (CIFs, green curves) and CIFs predicted from the same neurons’ spike phase distributions (purple curves). A) Stimulation at low frequencies (8-11 HZ), B) Stimulation at frequencies near neurons’ firing rates (38-55 HZ), C) Stimulation at high frequencies (79-86 HZ). The 4 rows are various examples for each of the 3 frequency ranges in no particular order. D) Goodness-of-fit evaluated by Pearson correlations between predicted and empirical CIFs across all GPe neuronal pairs in the sample as a function of driving frequency. E E) Scatter plot of amplitudes of CIFs that were predicted from the PDFs vs. the amplitude of the empirical CIFs (dashed line is the identity line).

What is striking about the various pairwise CIFs (Figs 4A-C) is that they exhibit diverse delays between the neurons, that depend on their iPRCs, firing rates and stimulus frequencies. This occurs even though each neuron was recorded separately on different brain slices in different mice, and neurons are therefore entirely independent of each other.

In summary, our analysis demonstrates that the iPRCs beget the return maps, which beget the phase distributions, which beget the pairwise CIFs. In other words, the heterogeneity in iPRCs is sufficient to explain the diversity of pairwise CIFs. Of particular relevance to previous interpretations of GPe neuron cross-correlations, our analysis shows that neurons driven by common input are not generally expected to produce CIFs with zero phase delay.

The two main features of CIFs of pairs of neurons subjected to common drive are their amplitude and their phase delay (or just “phase”). The amplitude indicates the strength of association between the neurons. In order to characterize how the amplitude of the CIFs calculated for the pairs of independent, sinusoidally-driven GPe neurons depends on the driving frequency, we plotted the maximal amplitude of the pairwise CIFs as a function of two parameters: the driving frequency and the geometric mean of the intrinsic firing rates of the two neurons (Fig. 5A). The first salient feature of this rendition is the large amplitudes at high driving frequencies particularly for the pairs with the lower intrinsic firing rates. The reason for this is that when the driving frequency is high, all spiking for all neurons tends to occur at the peaks of the stimulus that is closest to the end of the intrinsic period of the neurons (e.g., Fig. 1A, bottom). On the one hand, this input does not entrain the much slower neuron. Nevertheless, the rapid upswing closest to the end of the intrinsic period will recruit the next spike, and so the spiking for both cells occurs tightly around the phase of the peak (i.e., 0.25) of the stimulus cycle (Fig. 4C). The peak of the CIF will be high because it is the cross-correlation function of two unimodal, narrow distributions located around phase 0.25 of the input. The second salient property of the surface plot in Fig. 5A is the ridge of large amplitudes that runs diagonally along the curve where the driving frequency equals the geometric mean firing rate. This is intuitively clear, because it represents the case where one or both of the GPe neurons are being driven near their intrinsic firing rate. This too will give rise to narrow, unimodal phase distributions, which in turn will contribute to a prominent peak in the CIF.

**Figure 5:**
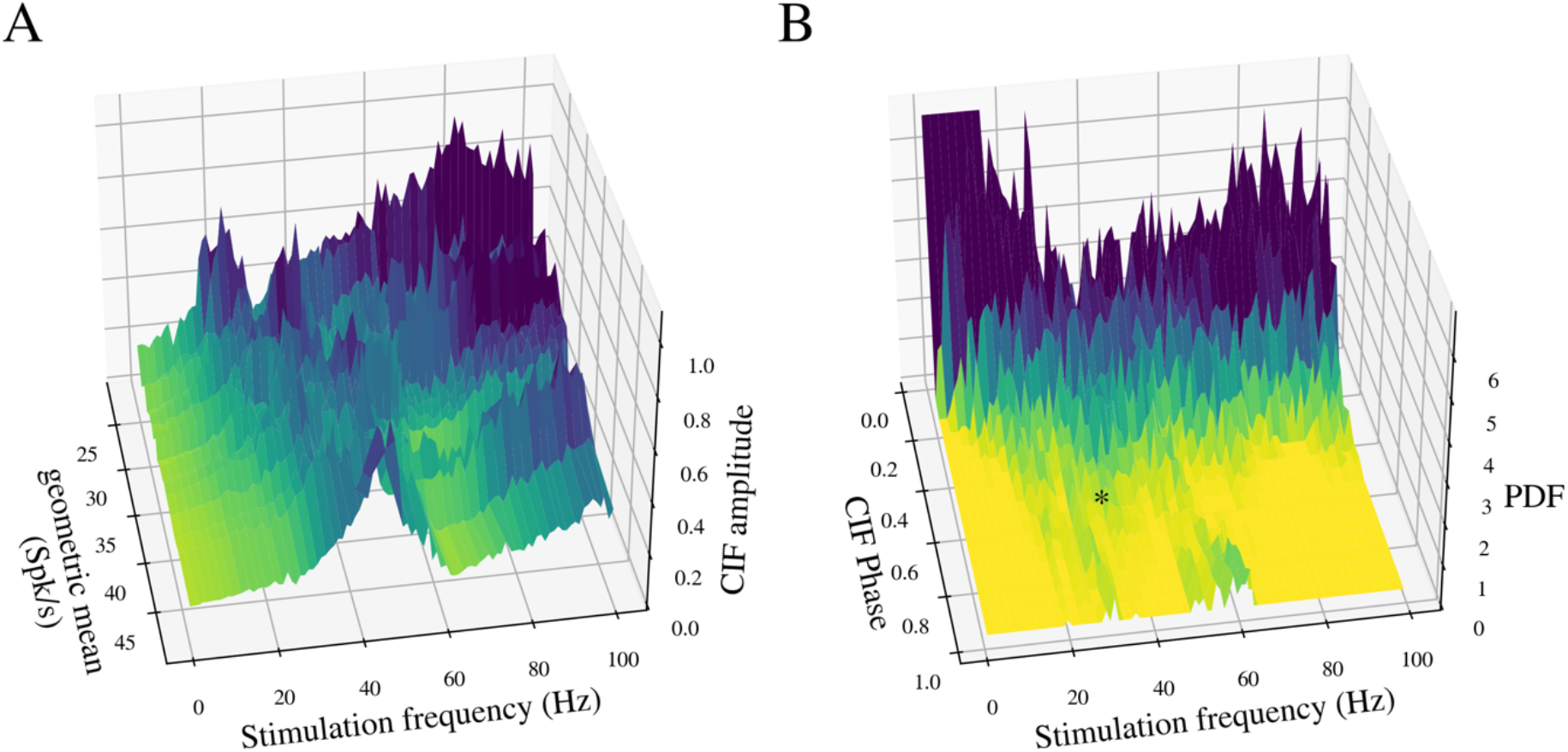
Experimental CIFs show frequency-dependent amplitude and broad distributions of phase delays. A) Amplitude of CIF as a function of two parameters: the driving frequency and the geometric mean of the intrinsic firing rates of the two neurons. B) Probability distribution function (PDF) of the CIFs’ phase delays across all pairs of GPe neurons as a function of the driving frequency, asterisk denotes occurrence of antiphase CIFs in the beta range. These data show that pairs of GPe neurons had a broad distribution of phase delays, particularly in response to driving frequencies in the beta range.

To characterize how the diversity of phase delays between neurons depends on the driving frequency, we plotted the probability distribution functions (PDFs) of the CIFs’ phase delays across all pairs of GPe neurons as a function of the driving frequency (Fig. 5B). This plot reveals that the broadest distribution of phase differences occurs for frequencies near the intrinsic firing rates of the GPe neurons (approximately 30 spikes/s) and at roughly double that frequency, which are the frequencies at which the strongest phase locking occurs. Note that in Fig. 5B there is a truncated large and narrow peak located at very low driving frequencies and at zero phase delay. At these very low driving frequencies the spiking of both neurons is rate modulated (*e*.*g*., Fig. 1A, top), which means that the CIF is a cross-correlations of the *rates* of two GPe driven by an identical sinusoidal input, so naturally both rates will peak at the same phase and the cross-correlation function will always peak at zero.

This analysis also demonstrates that beta-frequency periodic driving of completely independent GPe neurons is expected to generate CIFs with a broad distribution of non-zero phase delays.

### Intranuclear connectivity decorrelates entrained GPe neurons without changing phase delays

We found that unconnected GPe neurons, driven by the same stimulus frequency, exhibit broad distributions of phase delays in their CIFs, and the amplitudes of these CIFs depend on the cells’ firing rates and the stimulus frequency. To investigate the impact of local connectivity within the GPe on the CIFs of neurons receiving common sinusoidal inputs, we employed a network of simulated phase model GPe neurons.

As a preliminary step, we examined whether unconnected phase model neurons could replicate the behavior observed in uncoupled neurons in our experiments. Similar to the experimental data, unconnected phase model neurons were subjected to sinusoidal inputs ranging from 1 to 100 Hz. The stimuli were presented to a set of 1000 GPe model neurons with iPRCs and firing rates comparable to those in the experimental sample. We applied the same CIF analysis used for the experimental data to the resulting spike trains (see Fig. 6).

**Figure 6:**
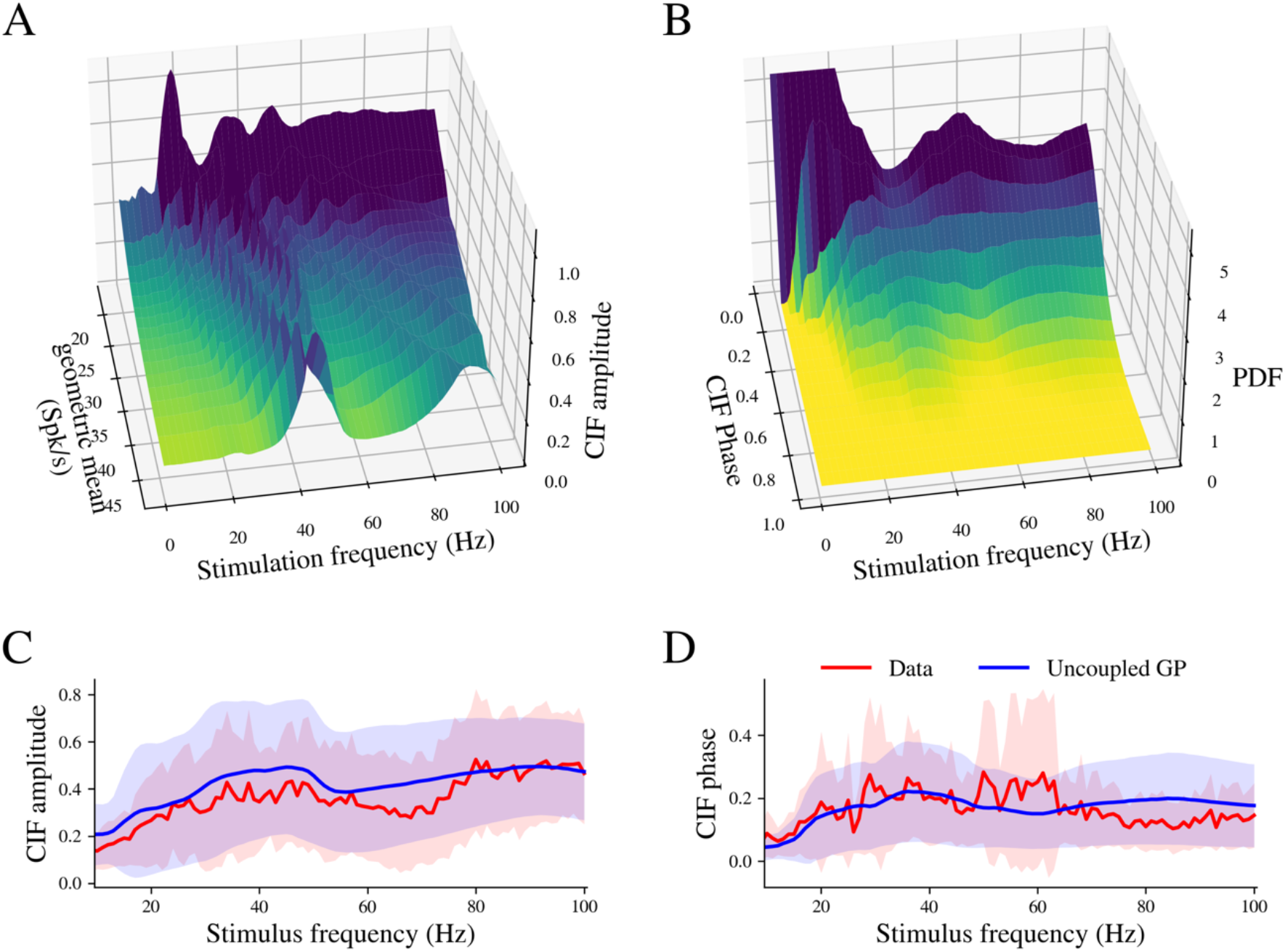
Unconnected *in silico* phase model neurons produce CIF amplitude and phase delay distributions that closely resemble the experimental data. A) Amplitude of CIF as a function of two parameters: the driving frequency and the geometric mean of the intrinsic firing rates of the two neurons. B) Probability distribution function (PDF) of the CIFs’ phase delays across all pairs of GPe neurons in the unconnected network as a function of the driving frequency. Mean *±* SD of CIF amplitude (C) and angle (D) as a function of stimulation frequency for experimental data (red) and unconnected model neurons (blue).

The amplitude of the pairwise CIFs in unconnected phase model neurons was reminiscent of the profile observed in the experimental data (compare Fig. 5A and Fig. 6A), including the ridge of large amplitudes running diagonally on the stimulation frequency versus pairwise geometric mean plane. The mean and standard deviation of CIF amplitudes for every stimulation frequency in the set of unconnected phase model neurons were similar to those observed in the experimental data (see Fig. 6C). Furthermore, the CIF phase delay angles across the stimulus frequencies for unconnected phase model neurons closely matched the distributions observed in the experimental data (compare Fig. 5B and Fig. 6B).

Next, we utilized the network model to study the impact of neuronal connectivity on CIFs. When connected, GPe neurons constantly receive a barrage of synaptic inputs from other GPe neurons. This barrage of inhibitory synaptic inputs decreases the regularity of inter-spike intervals and thereby alters the response to external inputs (Simmons et al., 2020). To quantify the specific contribution of local connectivity, we compared the CIFs from the connected network with those from an equivalent ensemble of unconnected phase model neurons, and a separate ensemble of unconnected neurons receiving a steady inhibitory barrage that approximated the average inhibition of the connected network but without its correlated temporal structure. To facilitate a direct comparison of CIFs, a constant depolarizing current was applied in both scenarios to offset the inhibition-induced reduction in firing rate. The same set of phase model neurons was used in all three configurations.

The principal effect of local connectivity on neurons’ CIFs is a dramatic decrease in CIF amplitude across all frequencies of stimulation studied, as shown in Fig. 7A. This reduction can be partly attributed to the irregularity induced by the synaptic barrage (Fig. 7B). To clarify this, we plotted the distribution of CIF amplitudes for all pairs in the sample [*N×(N-1)/2* pairs, *N = 1000*] at three stimulation frequencies (8, 35, and 80 Hz, Fig. 7C). Here, connected neurons (green) display a distribution skewed toward lower CIF amplitudes at all frequencies compared to unconnected neurons (black), while the unconnected neurons with barrage (red) show a similar skew, but only at the high frequency of 80 Hz. The influence of synaptic coupling on CIF phase delays is subtler (Fig. 7D); connected neurons and those receiving the barrage (green and red, respectively) exhibit distributions that are slightly skewed toward in-phase angles compared to unconnected neurons (black). It is worth mentioning the bimodal distribution of CIF phase delays at 35 Hz in neurons receiving the inhibitory barrage (red line 7D, middle panel). The origin of this distribution is explained by the neurons’ firing rates. The peak near zero delay is populated by neurons pairs whose rates are both above or below the stimulation frequency. The peak near 0.3 is populated by neurons pairs whose rates are one above and one below the stimulation frequency. The bimodal distribution is also present in connected and unconnected neurons (green and black lines) but with a greater overlap. The effect of CIF amplitude and subtler effect on CIF phase are also evident in the examples shown in Figs. 7E1 and E2, where we present CIFs of two neuron pairs at different stimulation frequencies. It is evident that the connected neurons (green) exhibit a reduced CIF amplitude at all frequencies, compared to the unconnected neurons (black), and this is true for the unconnected barrage configuration (red) only at the high stimulation frequency of 80 Hz. The CIF phase delay angle remained relatively stable across configurations but tended to gravitate toward antiphase angles as the stimulation frequency was increased.

**Figure 7:**
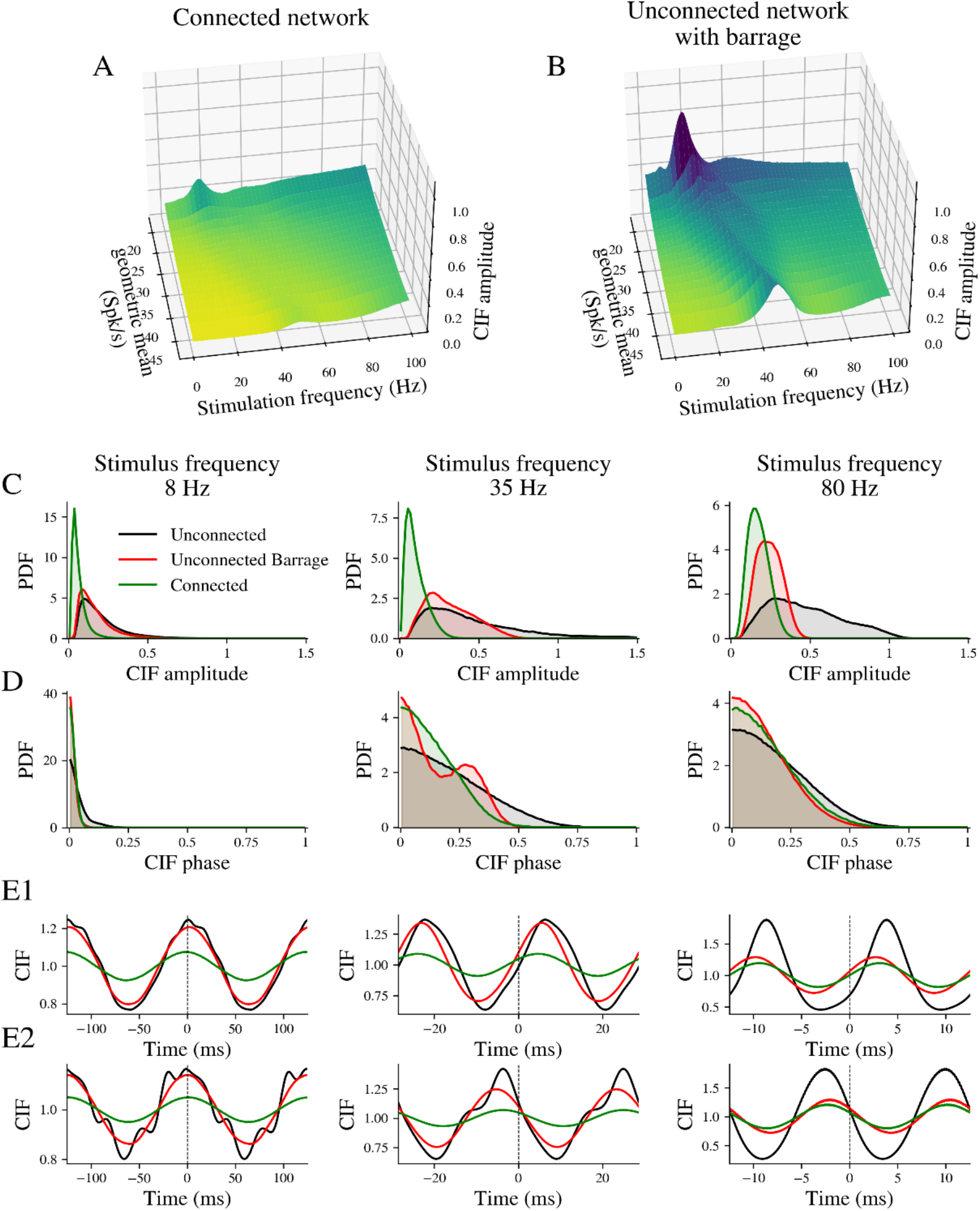
Impact of local neuronal connectivity on CIFs in GPe neurons. CIF amplitude as a function of stimulation frequency and the geometric mean rate of the neuron pair in the connected network (A) and for the unconnected network with the inhibitory barrage (B). C) Probability density functions of CIF amplitudes at selected stimulation frequencies (8 Hz, 35 Hz, and 80 Hz). Black, red, and green lines represent unconnected neurons, unconnected neurons with barrage, and connected neurons, respectively. D) Probability density functions of CIF angles at same stimulation frequencies as in C. E1 & E2) Example CIFs from two pairs of neurons at different stimulation frequencies for each model configuration.

These results were not dependent on the specific structure of the iPRCs in the GPe network, because repeating the same simulations with an identical iPRC for each neuron resulted in CIFs with similar amplitudes and phases (not shown). Moreover, altering the network architecture from the small-world to random or rich-club organization (with a long tail in the distribution of postsynaptic connections per neuron) while maintaining the mean number of connections, did not significantly alter the distribution of CIFs amplitudes and phases. Changing the total number of connections did impact CIFs amplitudes, but not phases. As connectivity decreased CIFs amplitudes smoothly approached the disconnected distribution of amplitudes, while increasing the number of connections decreased the CIFs amplitudes (not shown). Thus, the attenuation of CIF amplitudes by local GPe connectivity is robust to changes in the intrinsic properties of the GPe neurons that alter their iPRCs and to changes in network configuration. These findings suggest that local synaptic connections in the GPe decorrelates GPe neurons when they are driven by common input, by reducing CIF amplitudes without notably altering their phase delays.

## Discussion

### Phase models of GPe neurons fully predict their entrainment to oscillatory drive

We combined experiments and modeling to investigate the phase delays among GPe neurons subjected to a common oscillatory drive. We first addressed the question of how oscillatory drive determines the nature of the entrainment of an individual GPe neuron. For this, we recorded mouse GPe neurons in the perforated patch configuration while driving them with oscillatory current injections in the frequency range of 1-100 Hz. As described previously (Wilson, 2017; Tiroshi and Goldberg, 2019), we found that as the driving frequency approaches the neuron’s intrinsic frequency the neuron tracks the driving frequency. This tracking is accompanied by an increase in the VS, which reflects a sharpening of the spike phase distributions. The present study adds to these previous studies by revealing that the shape of the neuron’s iPRC determines the particular stimulus phase around which the spike phase distribution centers. In particular, by measuring the iPRC of each of the GPe neurons that were subjected to the oscillatory driving, we were able to show that the phase at which a neuron phase locks to its input, when driven at or around its intrinsic firing rate, is determined primarily by the angle of the fundamental Fourier mode of the iPRC. More generally, we found that by integrating the neuron’s phase equation (which takes the neu’on’s intrinsic firing rate and its iPRC into account) to produce a return map, then with appropriate additive noise, we were able to reconstruct the spike phase distribution at all frequencies, even in the absence of entrainment.

### The phase model of GPe neurons explains their temporal correlations in the presence of common oscillatory drive

After showing how the spike phase distribution can be derived from the individual neurons’ iPRC and phase equation, we showed that the structure of the CIF estimated for a pair of independent GPe neurons driven by a common oscillatory drive can be derived from the cross-correlation function of their corresponding spike phase distributions. Importantly, our analyses show that the phase delays of these CIFs can distribute broadly, particularly when the neurons are driven at frequencies approaching their intrinsic firing rates. To see why this is, note that the cross-correlation function of two distributions of random variables is essentially the distribution of the difference between these random variables. Therefore, the mode of the distribution of the CIF phase delays of a given pair of neurons, will be located at the difference between the modes of their respective spike phase distributions. We saw that as the driving frequency approached the intrinsic firing rate of a GPe neuron there was an increase in the stimulus phase at which the neuron phase locked (VS angle in Fig. 1E). Consequently, when one of two GPe neurons becomes entrained the differences between the modes of their spike phase distributions will become larger.

### Local connections between GPe neurons decorrelate them

The conclusion from these experiments is that connectivity among GPe neurons is not necessary to account for the broad distribution of phase differences among these cells seen in their CIFs, for example, in parkinsonian primates (Nini et al., 1995; Raz et al., 2000). We therefore asked, how would connectivity among GPe neurons affect the structure of their pairwise CIFs. To this end, we used an *in silico* approach, and found that when modeling the known connectivity between GPe neurons, the amplitudes of the pairwise CIFs were reduced. This finding agrees with previous studies of ours that have shown the local connectivity among GPe neurons serves to reduce the strength of their pairwise correlations, or “decorrelate” them (Wilson 2013; Olivares et al., 2022). It is not clear whether GPe collaterals are the primary influence that decorrelates GPe neurons, or whether their lack of pairwise correlation *in vivo* (Nini et al., 1995; Raz et al., 2000) is caused by the absence of overlap in joint input that neighboring GPe neurons receive (Wilson, 2013). Regardless, our simulation study suggests that even when (model) GPe neurons receive common oscillatory input, their local connectivity serves to decorrelate them. Importantly, this *in silico* result was robust to a battery of variations we applied to the iPRCs used in the simulation, indicating that decorrelation is a general outcome of the presence of local GPe connectivity.

An additional conclusion from this study is that in contrast to the amplitudes of the CIFs, which are attenuated by the collaterals, the phase delays among GPe neurons are essentially unaffected by the local connectivity. Thus, we can summarize that, while the amplitudes of the GPe CIFs are controlled by their local connectivity, the phase differences result from the intrinsic properties of the individual GPe neurons, which shape their respective iPRCs.

### Consistent phase relationships among neurons do not imply a direct interaction between them

There is a large body of work on phase relationships among GPe neurons (and other basal ganglia neurons) during oscillatory brain activity, whether they arise pathologically, as in various models of parkinsonism where beta oscillations dominate (Nini et al., 1995; Raz et al., 2000; Raz et al., 2001; Goldberg et al., 2004; Mallet et al., 2008) or under more physiological conditions such as sleep (Salih et al., 2009; Mizrahi-Kliger et al., 2018) and anesthesia (Magill et al., 2004; Magill et al., 2006; Slovik et al., 2017). In these studies, while acknowledging that correlation does not imply causation, finding consistent phase relationships among various neurons or neuronal populations was often interpreted as an indication of some direct or indirect interaction or influence between these neurons or populations (Perkel at al., 1967). For example, if two populations are anti-phase, then one inhibits the other (Goldberg et al., 2003; Magill et al., 2004; Magill et al., 2006; Mallet et al., 2008; Mallet et al., 2012, Abdi et al., 2015). Our findings provide a clear counterexample to this line of reasoning. In our data there is no interaction between the various GPe neurons recorded (because they are recorded separately on different brain slices from various animals), yet when subjecting them to the same oscillatory drive they give rise to a rich slew of calculated CIFs with various amplitudes and phase delays spanning all the way from in-phase to anti-phase (in a frequency dependent manner). Thus, it is important to consider that when a particular postsynaptic neuron exhibits a consistent phase with respect to some global oscillatory input, this phase may be best explained by the intrinsic properties of the neuron, as captured by its iPRC, which dictate the phase at which it entrains. The fact that each GPe neuron responds to a common input at a phase that is determined by its individual iPRC, implies that their output can remain phase incoherent with respect to each other even when they are subjected to a coherent input.

### The spectral decomposition of the iPRC

Our study underscores the spectral decomposition of the iPRC to its Fourier modes as an important property that elucidates how the iPRC interacts with oscillatory input. While the full solution of the phase model is needed in order to produce a return map, which in turn determines the precise phase at which locking occurs, we found mathematically that this phase is very nearly the angle of the fundamental Fourier mode of the iPRC. This means that two iPRCs that share the same fundamental mode will phase lock at roughly the same phase of the oscillatory drive.

iPRCs are typically thought of (and defined as) a way to characterize the responsivity of a neuron to a brief perturbation occurring at specific phase of the neuron’s firing cycle. Until recently it was also usually measured that way empirically (Reyes and Fetz, 1993). However, iPRCs also dictate how the phase of the oscillator will be impacted by extended perturbations that span the entire cycle (or more) by integrating the product of the input and the iPRC. In this case, because the oscillator and the iPRC are both periodic, the natural way to analyze the system is through its Fourier spectral decomposition (Goldberg 2013). In our case, this analysis demonstrates that it is these modes that determine the properties of the entrainment.

### Relevance to beta oscillations in parkinsonian primates

Our study may shed light on the structure of the CIFs observed among GP neurons in parkinsonian primates (Nini et al., 1995; Raz et al., 2000). It suggests that the oscillatory CIFs exhibited in these animals are the result of a global beta oscillation that entrains GPe neurons. It also suggests that the appearance of oscillatory CIFs does not require a change in the connectivity among GPe neurons. Moreover, the broad distribution of phase delays observed in the CIFs of GPe neurons in parkinsonian primates may reflect the intrinsic properties of these neurons with little bearing on the coupling between them. It should be mentioned that in our experiments the frequency ranges in which the CIFs exhibited the broadest distribution of phase delays (calculated for completely independent neurons) were close to the intrinsic firing rates of the GPe neurons, which were in the 10-40 spikes/s range. In primate GPe, the firing rates may be considerably higher than the beta frequency in which the pairwise CIFs of GPe neurons from parkinsonian primates exhibit a broad distribution of phase delay angles. One way to determine whether our explanation for the broad distribution of CIF delays applies to the broad distribution of phase delays observed in parkinsonian primates would be to estimate the global oscillatory signal present in those animals, for example, through recording of the local field potentials (LFPs) and constructing the spike phase distribution with respect to the phases of the oscillatory LFP. One could then test whether the convolution of these distributions gives rise to the empirical CIFs, which is reminiscent of a previous study showing the GPe CIFs in parkinsonian primates can be predicted from the LFP (Goldberg et al., 2004).

## Acknowledgments

This work was supported by National Institute of Neurological Disorders and Stroke Grant R35NS097185 to C.J.W. The authors would like to thank Rostislav Likhotvorik for his excellent technical support, and Drs. Lior Tiroshi, Matthew Higgs and Hitoshi Kita for helpful advice and comments on the manuscript. J.A.G. was on sabbatical at UTSA during the performance of this work.

## Notes

**Conflict of Interest Statement** The authors declare no competing financial interests.

### Competing Interest Statement

The authors have declared no competing interest.

### Summary of Updates

Revision includes a new introduction, more mathematical exposition, additional simulations, and clearer figures.

